# Non-linearity in motor unit velocity twitch dynamics: Implications for ultrafast ultrasound source separation

**DOI:** 10.1101/2023.03.24.533983

**Authors:** Emma Lubel, Bruno Grandi Sgambato, Robin Rohlén, Jaime Ibáñez, Deren Y Barsakcioglu, Meng-Xing Tang, Dario Farina

**Affiliations:** Department of Bioengineering, Imperial College, London, United Kingdom; I3A, University of Zaragoza, Spain; Department of Biomedical Engineering, Lund University, Sweden

**Keywords:** B-mode, Intramuscular Electromyography, Motor Units, Surface Electromyography, Ultrasound

## Abstract

Ultrasound (US) muscle image series can be used for peripheral human-machine interfacing based on global features, or even on the decomposition of US images into the contributions of individual motor units (MUs). With respect to state-of-the-art surface electromyography (sEMG), US provides higher spatial resolution and deeper penetration depth. However, the accuracy of current methods for direct US decomposition, even at low forces, is relatively poor. These methods are based on linear mathematical models of the contributions of MUs to US images. Here, we test the hypothesis of linearity by comparing the average velocity twitch profiles of MUs when varying the number of other concomitantly active units. We observe that the velocity twitch profile has a decreasing peak-to-peak amplitude when tracking the same target motor unit at progressively increasing contraction force levels, thus with an increasing number of concomitantly active units. This observation indicates non-linear factors in the generation model. Furthermore, we directly studied the impact of one MU on a neighboring MU, finding that the effect of one source on the other is not symmetrical and may be related to unit size. We conclude that a linear approximation is limiting the decomposition methods to decompose full velocity twitch trains from velocity images, highlighting the need for more advanced models and methods for US decomposition than those currently employed.

## 1. Introduction

IN voluntary movements, action potentials (APs) discharged by motor neurons (MNs) reach muscle fibers, eliciting their contraction. The MN and innervated fibers constitute the motor unit (MU) – the smallest functional unit of human movement. At present, tools have been developed which allow us to decompose the spiking activity of MUs, which is a useful resource for decoding neural activity in humans. Most commonly, high-density surface EMG (HDsEMG) is used for investigating MUs [1]–[3]. However, although HDsEMG is non-invasive, it has low spatial resolution [4]–[6] and poor penetration [6], [7], so only superficial muscle layers contribute to the signal. Intramuscular EMG (iEMG) can measure the MU activity in deeper muscle layers, but it is invasive and can only detect a small area around the recording electrode sites [8]. Alternatively, ultrasound (US) has greater penetration and spatial resolution while still being non-invasive. Thus, it could provide a more complete picture of muscular activity than EMG-based approaches.

In contrast to EMG, which relies on identifying the unique electrical APs generated by each MU, US-based MU decomposition identifies the times of occurrence of mechanical velocity twitches generated by each MU. This is possible using ultrafast US (> 1000 frames/s) [9], which has a sufficient temporal and spatial resolution to capture the mechanical displacement of muscle units. The times of mechanical velocity twitches generated by muscle units have a precise temporal association with the discharge times of the innervating MN so that US decomposition can be used to unravel the neural activity traveling along MNs. In this way, a mechanical measurement provides a neural interface with the output layers of the spinal cord.

While the source mixing model for EMG signals is linear, the motion of a muscle unit and its summation with those of other units likely results in complex interactions. The structure of skeletal muscles is extremely complicated, with dense packing of fibers surrounded by connective tissue, bone, skin, and blood vessels. When a MU contracts and relaxes, it pushes and pulls on nearby fibers and connective tissue and sets up propagating waves within the muscle tissue [10], [11]. Overlapping MU territories present intermingled fibers from different MUs contracting at different times [12]. Furthermore, a twisting motion of contracting MUs has been identified in both stimulated [13] and voluntary [14] contractions. Hence spatially distant regions are seen to move in unison. Importantly, unlike EMG, which is made of electric potentials of short duration, the elicited motion of muscle fibers has a smaller bandwidth and, therefore, longer duration.

Despite the complicated motion, methods have been proposed for the estimation of the neural drive to muscles (individual MU discharge times) via the decomposition of US image series assuming linear models [15]–[17]. These methods treat the problem as a blind source separation challenge, where each MU is the source of a unique spatio-temporal signal. To solve the separation problem, spatio-temporal independent component analysis (st-ICA) has been applied [15], [18]. The main assumptions of this approach are that the source signals are independent of each other and are all non-gaussian and that they are mixed following a linear instantaneous mixing model. Using these methods at low force levels (such that a single unit was detected using EMG), approximately 30% of MUs detected using needle EMG (gold standard) were also detected from US, with a rate of agreement (RoA) in the identified discharge times of approximately 75% [16], [19]. This performance level is substantially poorer than observed when decomposing US signals simulated by linear superposition models (75-95% of the MUs detected with a RoA of 90% for up to 20 active MUs [15]). This discrepancy may point out that the linear instantaneous model assumption for the sources is being violated and, therefore, the method fails to properly identify the sources. Further understanding the intricacies of any non-linearities involved and the limits of a linear approximation is key for developing future US decomposition algorithms, which would enable full decoding of the neural drive to the entire muscle cross-section, something impossible with current EMG methods.

The main objective of this study was to determine if the superposition of individual MU activity can reasonably be considered linear from a velocity field perspective. For this purpose, we analyzed if the velocity twitch profile of a MU is influenced by the activity of other MUs. We used a combination of HDsEMG and US with a modified spike-triggered averaging (STA) method to obtain the response in a given region to a MU firing, termed the velocity twitch profile. We compared the velocity twitch profiles of the same MUs during voluntary isometric contractions at 2%, 5%, 10%, and 20% of the maximum voluntary contraction (MVC) force. Further, in a second study, we used a combination of iEMG and US to isolate extremely close muscle units and to study their influence on each other in terms of US images.

### II. METHODS

Two experiments were conducted. Experiment 1 combined HDsEMG and ultrafast US, which allowed for large populations of MUs to be analyzed. Experiment 2 used iEMG and ultrafast US to detect MUs clustered in a small region, allowing for a more specific analysis of MU interactions. The methods section will be divided into two sections describing each experiment.

#### Experiment 1: HDsEMG and US

#### 1) Participants

12 healthy participants were recruited for this study. However, system and synchronization errors caused data from two participants to be unusable. Hence data from 10 participants were used for all the analyses (*n* = 10, 26.2 ± 2.9 yr, 173.2 ± 7.3 cm, 69.1 ± 12.0 kg). Before the experiment, the experimental protocol was clearly explained to the participants, and they signed an informed consent form. Procedures and experiments were approved by the Imperial College Research Ethics Committee (ICREC reference: 20IC6422) in accordance with the Declaration of Helsinki.

#### 2) Instrumentation

This experiment required the synchronized acquisition of HDsEMG, ultrafast US, and force data. The HDsEMG signals were recorded using two grids of 64 electrodes each (5 columns and 13 rows; gold coated; 8 mm interelectrode distance; OT Bioelettronica, Torino, Italy). Signals were recorded in monopolar derivation, amplified, sampled at 2048 Hz, A/D converted to 16 bits with gain 150, and digitally bandpass filtered between 10 Hz and 500 Hz with an EMG pre-amp and a Quattrocento Amplifier (OT Bioelettronica, Torino, Italy). The US data were recorded using an L11-4v transducer with 128 elements and a center frequency of 7.24 MHz. The Vantage Research Ultrasound Platform (Verasonics Vantage 256, Kirkland, WA, USA) was controlled using custom codes written in MATLAB (Mathworks, Massachusetts, USA). Single angle plane wave imaging at 1000 frames per second was used, resulting in 30,000 data frames for 30 s recordings. Delay and sum beamforming produced B-mode images (357 by 128 pixels). For synchronization, the Verasonics trigger function was used to produce a 1-μs active low output, which was elongated by an Arduino Uno and fed into the Quattrocento Amplifier to allow for alignment of the recorded US and HDsEMG data. Finally, the force data were recorded using an ankle dynamometer and fed through a Forza force amplifier (OT Bioelettronica, Torino, Italy) into the Quattrocento Amplifier.

#### 3) Experimental procedures

The skin area above the tibialis anterior muscle (TA), chosen for its long muscle fibers and low pennation angle [20], was shaved and cleansed with a chemical abrasive and alcohol. Next, the muscle was palpated, and the HDsEMG electrode grids were attached to the skin. The first electrode grid was placed proximally, following the direction of the muscle fibers. The second electrode was placed distally, leaving a 1 cm gap between the electrodes. The electrodes were secured using Tegaderm Film Dressings and self-adhesive medical bandages. Next, using a custom-designed 3D-printed probe holder, the ultrasound probe was attached to the leg in the gap between the EMG grids with the imaging plane perpendicular to the muscle fibers. As such, the resultant images were cross-sectional views of the TA muscle. A water-based US gel improved the coupling between the probe and the leg. Next, the participant was seated at a comfortable distance from a computer screen, and their leg was secured into an ankle dynamometer with their foot at a 90-degree angle to their leg. Soft padding was placed around the leg to provide comfort and secure the leg in position.

Real-time force feedback from the ankle dynamometer was used for this experiment, and as such, the participant was guided by on screen force ramps. Initially, the participant was instructed to perform the strongest dorsiflexion contraction they could, and their MVC force was recorded. Following this measure, they performed four ramp contractions at 2%, 5%, 10%, and 20% MVC. The ramps were 50-s long, with 5-s rise and fall times and a 40-s plateau. Once the participant reached the plateau and the force was judged stable, the experimenter began a 30-s US recording. Between each ramp the participant rested for 60 s. The protocol is illustrated in Fig. 1.

**Fig. 1.**
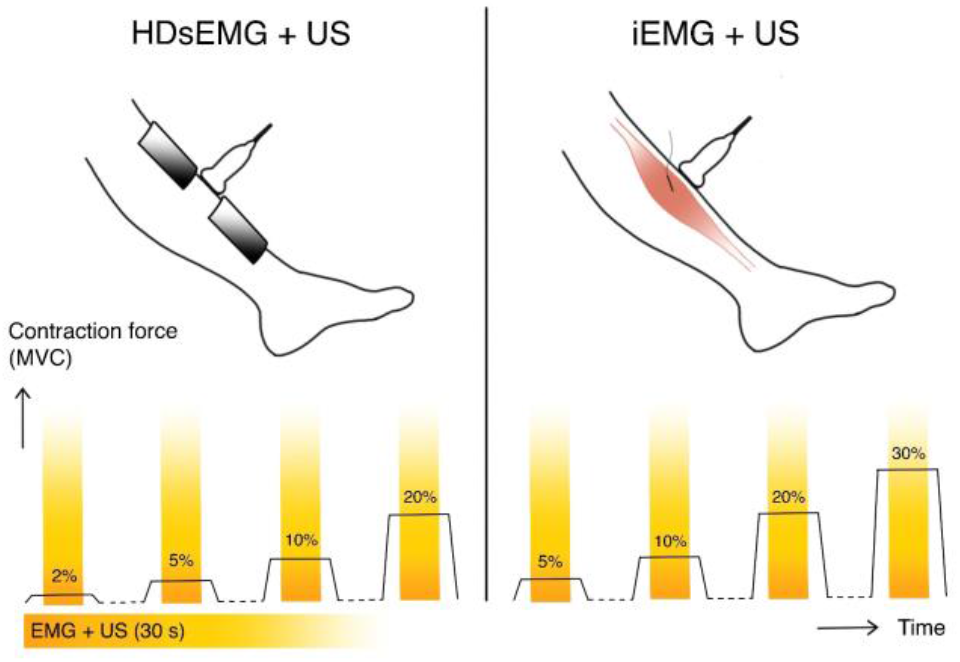
The experimental set-up and procedures for experiments 1 (left) and 2 (right). In both experiments ultrasound (US) was used, with the imaging plane perpendicular to the length of the muscle. Additionally, either high-density surface electromyography (HDsEMG) or intramuscular EMG (iEMG) was used. In each case, the participant followed force ramps by isometric ankle dorsiflexions with online feedback on the produced force, as shown at the bottom of the figure. At each force level plateau, a 30-s synchronised recording of US and EMG was taken.

#### 4) Data Processing

The HDsEMG signals were decomposed using a validated method [21], in four intervals, corresponding to the four contraction forces. On each interval, a fastICA algorithm was used to iteratively optimize a separation vector for each active MU, maximizing the sparseness of the sources and outputting an estimated discharge time series for each MU. The separation vectors from each interval were applied to all other intervals to identify the same MUs across different contraction levels. Then any duplicate MUs were removed. Hence the same decomposed unit was tracked across all intervals (and, therefore, forces). K-means clustering was used to separate the high peaks (corresponding to MU spikes) from the low peaks (other MU spiking activity and noise). The estimated discharge times were manually refined and edited by an expert [22]. MUs were retained if they were active in all four intervals or in all but the 2% MVC force interval. The US data was beamformed, and a 2D autocorrelation method [23] was used on the reconstructed radio-frequency data to calculate a tissue velocity series, using a sliding window of 2 ms in time and 1 mm in depth. In this work, a negative velocity indicated motion away from the probe (and hence the skin).

At each contraction level, a previously validated STA method [14], was used to detect MU motion domains. The processing and terminology are summarized in Fig. 2. In short, an STA of the velocity maps was computed, resulting in 100-frame (100 ms, 50 ms before, and 50 ms after firing time) videos for each MU. For each pixel in the image series, the squared sum of the STA divided by the variance of the curves making up the STA was calculated such that regions with high motion and low variability across triggers have high magnitudes. The map was then modulated with a -1 if the direction of the motion was away from the probe at the time of the EMG spike. The map was finally thresholded at 65% of the maximum (both towards and away from the probe). The resultant regions were termed the MU motion domains – the regions of the muscle cross section which moved synchronously in response to MU firings.

**Fig. 2.**
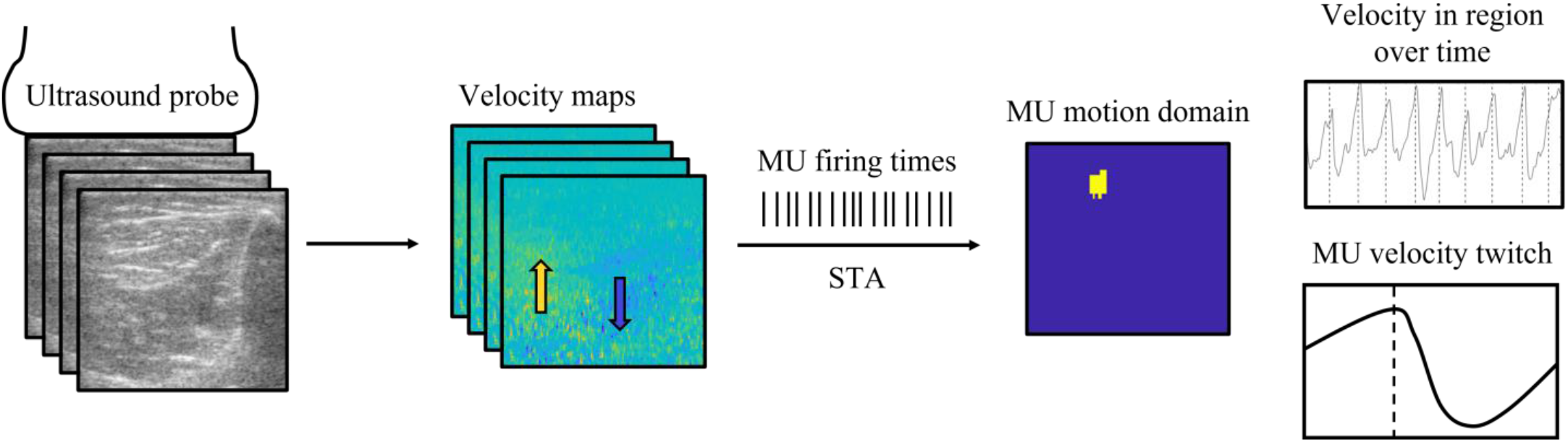
Diagram showing the processing pipeline for each contraction level. B-mode ultrasound images are recorded at 1000 frames per second and 2D autocorrelation methods are used to estimate the tissue velocity from the reconstructed radio-frequency images (either towards or away from the probe). Simultaneously recorded EMG signals were decomposed into motor unit (MU) firing times, then a spike-triggered averaging (STA) method was used to identify the MU motion domain (the region which moved synchronously in response to a MU firing), and the MU velocity twitch – the STA velocity profile in this region.

Although recordings were performed at 4 contraction levels, at 2% MVC force, very few MUs were identified. Since we tracked the same units across all contraction levels, low numbers of MUs obtained at 2% MVC would restrict our analysis to only 45 unique MUs (4.5 ± 4.1 per participant). Conversely, by excluding the 2% MVC force level from the analysis and considering only the 5%, 10%, and 20% contraction levels, 170 MUs (17 ± 12 per participant) could be identified and tracked across forces. Hence, most of our results come from analyzing three out of the four force levels.

Further processing was divided into two categories: processing on all four contraction levels and processing on just the highest three contraction levels. For the former, if a pixel was present in the MU motion domain in 3 or more of the 4 maps, it was selected as part of the region used for comparison. For the latter, if a pixel was present in the MU motion domain in 2 or more of the 3 relevant maps, it was selected as part of the region used for comparison. This ensured that the selected region was within the common MU motion domain. Some typical MU territories and common regions are shown in Fig. 3 for each scenario. MUs whose motion domains across contraction levels did not overlap were discarded. Hence the remaining MUs were able to be tracked with high spatial consistency across contraction levels. Next, the average STA within this set of selected pixels (i.e., the region selected for further analysis) was calculated for each force level. These were termed MU velocity twitches.

**Fig. 3.**
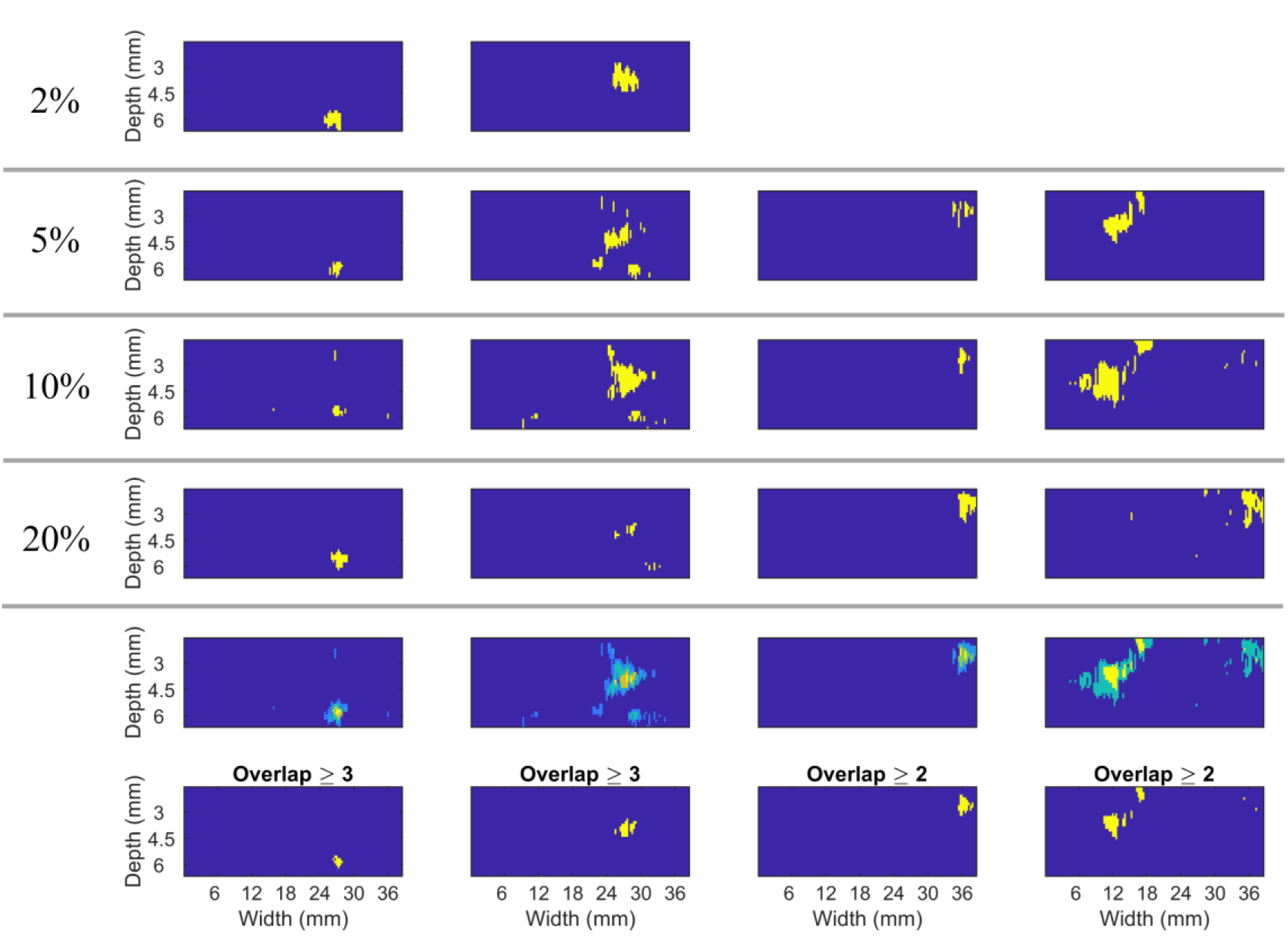
Four examples of motor unit (MU) motion domains at different contraction levels (yellow = MU domain), alongside their overlapping territories and the territories selected for analysis. Left two columns: MUs active at 2%, 5%, 10%, and 20% MVC (rows 1 to 4), their summed territories (row 5) and the region selected for analysis (row 6). The first column shows an example where there is little noise and the regions all clearly overlap. The second column shows an example where noise, especially at 20% MVC, results in a worse STA performance, hence overlap is only required in 3 or more maps. Right two columns: MUs active at 5%, 10%, and 20% MVC (rows 2 to 4), their summed territories (row 5) and the region selected for analysis (row 6). The third column shows an example where there is little noise and the regions all clearly overlap. The fourth column shows an example where noise, especially at 20% MVC, results in a worse STA performance, hence overlap is only required in 2 or more maps.

The MU velocity twitch peak-to-peak amplitude was compared across force levels. Furthermore, individual (non-averaged) velocity twitches were studied as follows: the correlation of each individual velocity twitch with the average MU velocity twitch across all force levels was calculated, and the twitches were sorted into either a ‘correlated’ group (correlation with the grand average twitch ≥ 0.5) or an uncorrelated group (correlation < 0.5). Given that correlation is dependent on the shape and not on amplitude, this criterion separated twitches into those with similar shapes (invariant of their peak-to-peak amplitude) and those with dissimilar shapes. One STA was then calculated for the correlated group and one for the uncorrelated group.

#### 5) Statistics

All results were statistically analyzed using the following steps. First, the normality of the distributions was tested using a Shapiro-Wilk test (p < 0.05). In all cases, at least one involved group failed the normality test. Thus, methods that do not require normality were used. In all cases, the data were paired (e.g., same MU at different contraction levels, same MU with different processing), so the non-parametric Friedman test was used, followed by a post-hoc Conover test with Bonferroni corrections for multiple comparisons. In these cases, the corrected p-value (weighted by the number of pairwise comparisons) is shown. The significance level was set as p < 0.05. The p-values are stated on each graph with 3 significant figures, and any p-values lower than 0.001 are labeled as p < 0.001.

### B. Experiment 2: iEMG and US

#### 1) Participants

One participant was recruited for this study (*n* = 1, 32 yr, 183 cm, 73 kg). Before the experiment, the experimental protocol was clearly explained to the participant, and they signed an informed consent form. Procedures and experiments were approved by the Imperial College Research Ethics Committee (ICREC reference: 19IC5641) in accordance with the declaration of Helsinki.

#### 2) Instrumentation

The instrumentation for recording the US, force, and synchronization data was the same as in experiment 1. Three thin-wire bipolar iEMG electrodes were used to record the EMG data (OT Bioelettronica, Torino, Italy). The acquisition of the EMG was performed using the Quattrocento amplifier (OT Bioelettronica, Torino, Italy) at a sampling frequency of 10240 Hz, bandpass filtered between 10 Hz and 4400 Hz.

#### 3) Experimental procedures

The skin above the participant’s TA was cleaned and wiped before the experiment began. Three thin wire bipolar EMG electrodes were inserted along the length of the muscle fibers at approximately 45 degrees until the tip of the insertion needle was approximately 1.5 cm deep. The first electrode was inserted to align with the center of the US probe. The insertion was done by an expert and guided using ultrasound to ensure the needle tip was positioned correctly. Once the electrode was correctly positioned, the needle was removed, leaving just the thin wire electrode in place. The US probe was attached to the leg as in experiment 1, and the leg was secured in the ankle dynamometer.

The participant performed an MVC contraction. Next, two repeats of four force ramps were performed at 5%, 10%, 20%, and 30% MVC. Each ramp consisted of a 5-s rise, a 40-s plateau and a 5-s fall. Once the participant reached the plateau, a 30-s US recording was stored (see Fig.1). Between each ramp, the participant rested for 60-s. Once one set of 8 ramps had been completed, a second wire electrode was inserted medially with respect to the first electrode, and the 8 ramps were repeated. Finally, a third electrode was inserted laterally with respect to the first electrode, and the 8 ramps were repeated again. This resulted in a total of 24 synchronized paired 30-s recordings of iEMG and US.

#### 4) Data Processing

The iEMG data were high-pass filtered at 1000 Hz and EMGLAB’s [24] automatic decomposition was used on the full iEMG signal. An expert visually corrected the spike trains for missing or double firings. The resultant templates and spike trains were exported to MATLAB, and any MUs with spike trains with inter-spike interval coefficient of variation greater than 30% were discarded [25]. The remaining MUs were used for further analysis.

The data processing for the US data was the same as in experiment 1. Using the velocity maps calculated from the US and the firing times from the iEMG decomposition, the same STA processing as described in experiment 1 was carried out, resulting in MU motion domain maps and velocity twitch STAs. Any MUs whose motion domains were not within the expected detection region of the EMG electrode or were noisy (scattered around the map rather than concentrated) were discarded. Next, pairs of nearby MUs were used for the analysis of how the activity of one MU (MU1) was affected by its neighbor (MU2) in two ways:

(i) For a given MU (MU1), the activity of a nearby MU (MU2) occasionally began mid-way through the contraction. Thus, MU1s activity could be separated into two time intervals: activity before the nearby unit was active and activity after. The *n* velocity twitches before MU2 was active were used to produce an STA representing the activity of MU1 in the ‘active alone’ condition. Then, *n* of the velocity twitches (selected randomly) in the second group (MU1 and MU2 both active) were used to produce an STA of MU1s activity in the ‘active together’ condition. This was repeated until the remaining number of twitches in the second group was less than *n*. The STAs were then averaged to produce an average twitch in the ‘active together’ condition. Finally, the STA of MU1 before and after MU2s activity began were compared to analyze the impact of MU2 on MU1.

(ii)The firing times of MU1 were categorized into two groups: those which aligned with firings from MU2 and those which did not. The window for alignment was adjusted on a case-by-case basis to ensure equal numbers in each group, always within the range of ±10-20 ms. Using each of these groups, an STA of the velocity profile in the motion domain of MU1 was calculated to produce an aligned spikes STA and an unaligned spikes STA. This enabled the study of quasi-individual MU activity versus activity alongside a nearby unit. The MUs were then investigated the other way around to determine if the impact of one MU on another is symmetrical.

## III. RESULTS

### A. Experiment 1: HDsEMG and US

On average, the peak-to-peak amplitude of the velocity twitch decreased as the contraction level increased. In other words, the average velocity twitch of individual MUs was suppressed as contraction force increased from 2% to 20% MVC. An example of this phenomenon is shown in Fig. 4A. The relative peak-to-peak amplitude of the velocity twitch profiles across the contraction levels is shown in Fig. 4B. For each contraction level, the distribution of peak-to-peak amplitudes was statistically significantly different (p < 0.05). This is shown in Fig. 5, where the 10% and 20% MVC contractions are compared with the 5% contraction, and the groups are statistically different (p < 0.05).

**Fig. 4.**
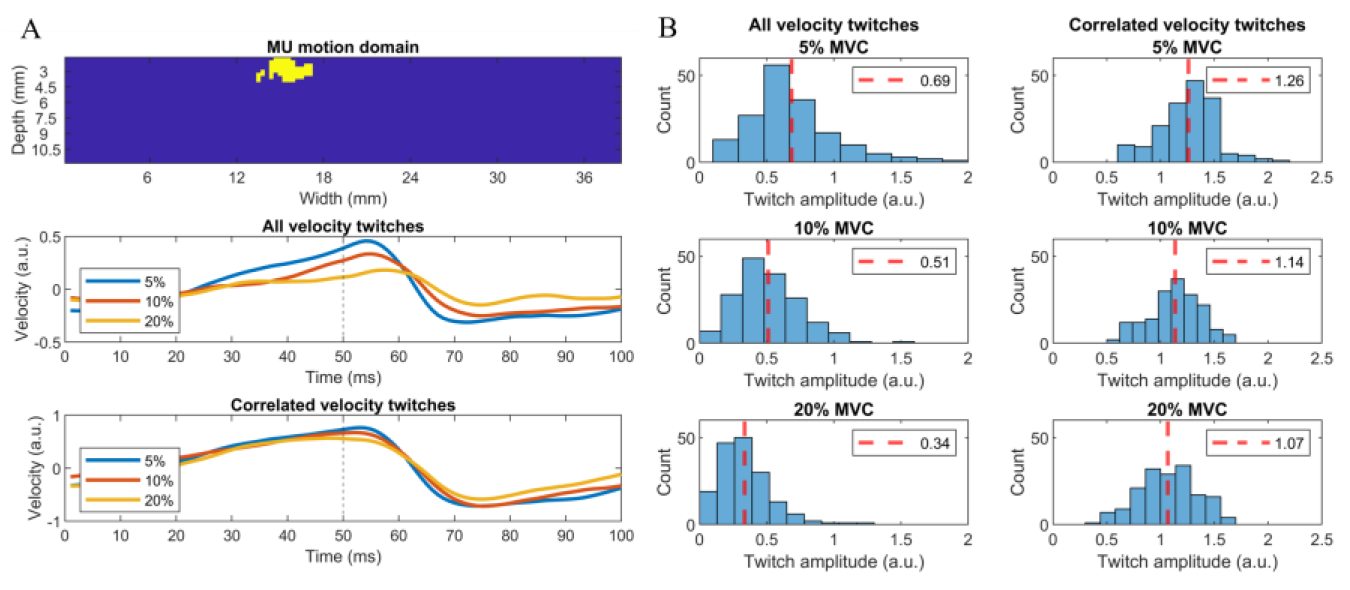
A: An example of a common region of the motor unit across contraction levels (top) and velocity twitch profiles, showing a decrease in peak-to-peak amplitude at increasing contraction levels (middle). When only the highly correlated twitches are included (bottom), the peak-to-peak amplitude is more consistent. B: Velocity twitch amplitude of spike-triggered average (STA) velocity twitch when all firings contribute to the STA (legt) decreases as the contraction level increases. All contraction levels are significantly different with p < 0.001. When only highly correlated velocity twitches were considered (correlation with overall average twitch profile ≥ 0.5) (right), the amplitude was more consistent across contraction levels. All groups were statistically different, with all p < 0.001, except between 10% and 20% where p = 0.0467 (with correction). In each case, the p-value was much higher than that for the left plots. In each plot the mean value is shown by the dotted red lines.

**Fig. 5.**
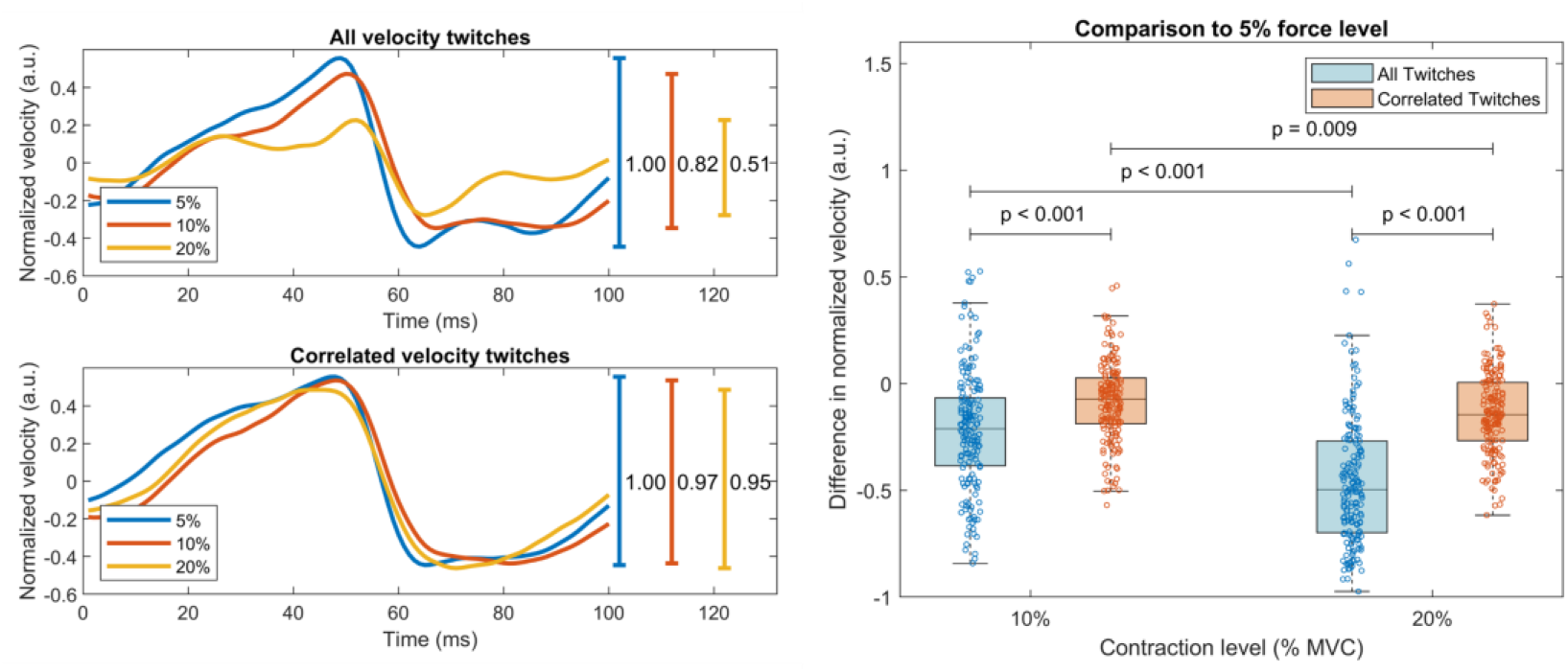
Left: Example of normalised velocity twitch sizes for the same MU considering all velocity twitches (top) and just correlated velocity twitches (bottom). Right: Peak-to-peak amplitudes of velocity twitch profiles at 10% and 20% MVC compared with 5% velocity twitch profile (i.e., the difference between the values marked on the right-hand side of the left plot). When all velocity twitches are used to produce the STA, the STA is suppressed for both 10% and 20% (p < 0.001, n = 170). When just the highly correlated twitches are considered, the 10% and 20% twitches are more similar to each other and those at a 5% contraction level (p = 0.009, n = 170).

It must be noted that, contrary to the global muscle force, the velocity twitch is a signal in space and time. At low contraction forces, it is possible to identify velocity twitches isolated from the others because of the sparseness of the signal in both space and time. In a series of velocity twitches, we assumed that some would be isolated while others were not. We identified the isolated velocity twitches by determining the group of twitches ‘correlated’ with the grand average STA (see example in Fig. 6). The correlated group comprises the twitches that are mostly preserved in individual activations and that, therefore, are least influenced by other MUs. When we used only the correlated velocity twitches to determine the STA, the peak-to-peak amplitude remained constant across contraction levels, as shown in Fig. 4 and Fig. 5. This is an interesting result since it indicates that when considering isolated twitches, the twitch response was not influenced by the force level or the number of active MUs. This result indicates that it is unlikely that the decrease in velocity twitch amplitude when considering the entire discharge time series is due to changes at the whole muscle level, such as stiffness (otherwise a decrease would also occur when selecting a group of twitches with correlated shapes). Rather, results reflect that the non-linear effects (causing the reduction in STA velocity twitch amplitude) are local and depend on the mechanical disturbance due to the activity of closely located MUs. With increasing MVC levels, we see a decrease in the percentage of twitches which fall into the highly correlated group (Fig. 7) due to an increase in the number of active nearby MUs.

**Fig. 6.**
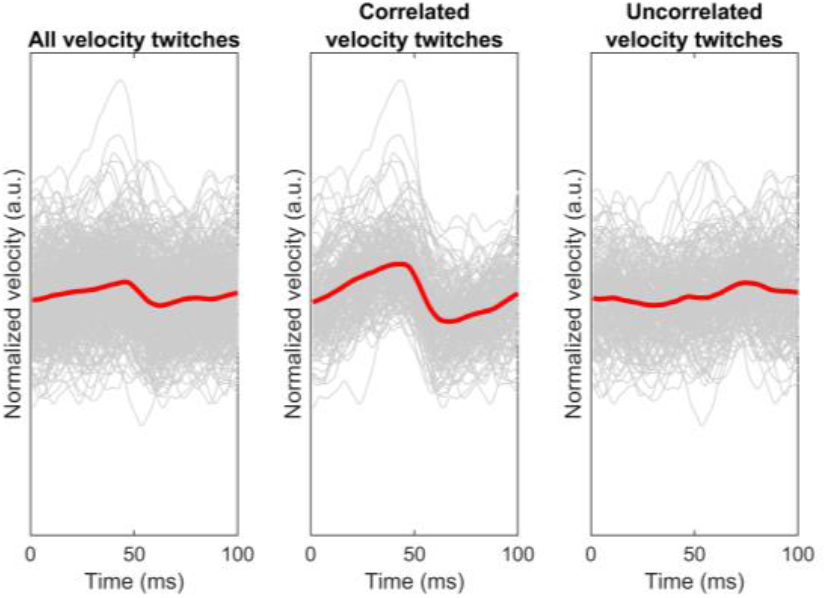
Plots to show effects of separating correlated and uncorrelated group for a 20% MVC contraction. Left: spike-triggered average (STA) produced using all velocity twitches. Middle: STA produced using velocity twitches with a correlation with the average twitch across all force levels of > 0.5. Right: STA produced using velocity twitches with a correlation < 0.5. With increasing force level, the size of the STA in the left plot decreases due to the number of contributing lines in the right plot increasing. However, the middle plot remains consistent – we are able to extract a ‘template’ twitch profile that is the same at all levels, but motion of nearby units causes an increasing number of twitches to not fit the expected velocity twitch pattern.

**Fig. 7.**
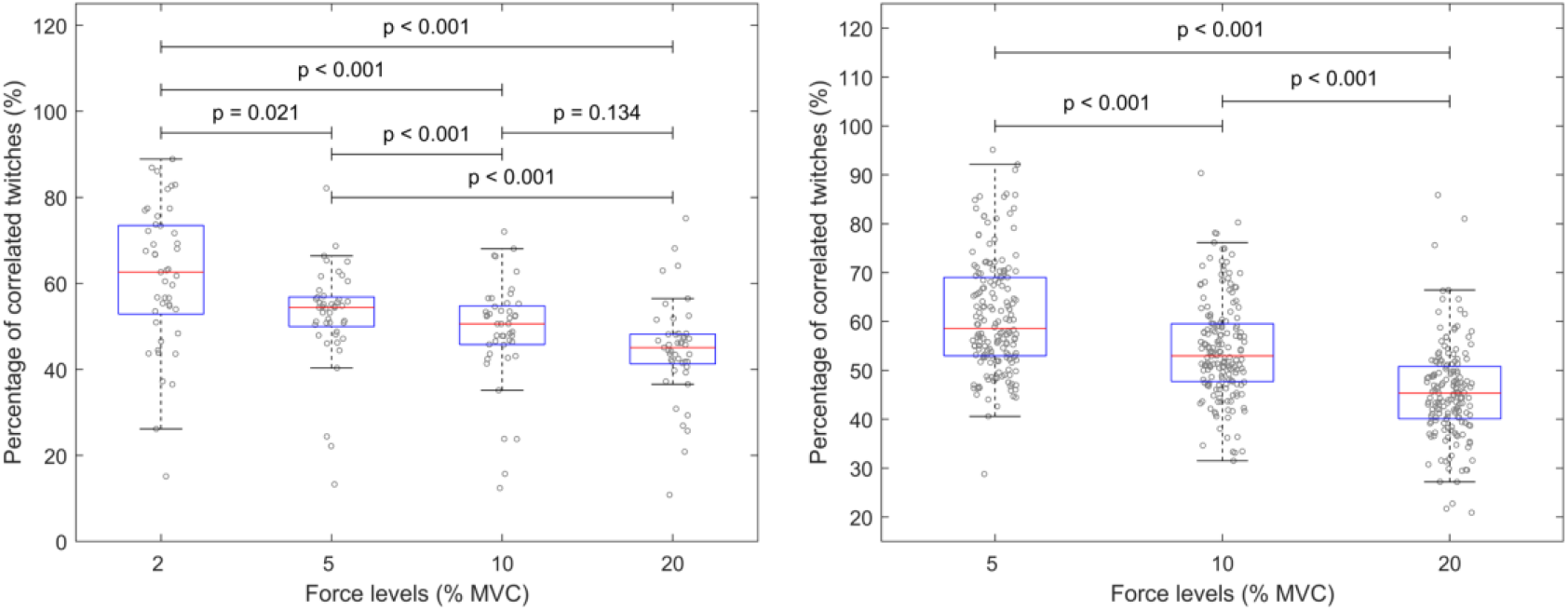
Boxplots showing percentage of velocity twitches which have a correlation ≥ 0.5 with the overall average velocity twitch STA. In each case, one dot at each force level corresponds to one motor unit (MU), so one MU will have a dot representing its percentage correlation at each force level. Left: For 2%, 5%, 10%, 20% MVC (n = 45, p-values have Bonferroni correction - multiplied by 6 for 6 pairwise comparisons, significance p<0.05). The number of correlated velocity twitches decreases with increasing contraction level, however 10% and 20% are not significantly different. Right: For 5%, 10%, 20% MVC (n = 170, p-values have Bonferroni correction - multiplied by 3 for 3 pairwise comparisons, significance p<0.05). Here the group sizes are much larger, and all groups are significantly different.

### B. Experiment 2: iEMG and US

In this experiment, MUs close to each other were identified due to the small detection volume of the bipolar iEMG electrodes. In a number of recording intervals, the activity of one unit began after the beginning of the recording. This allowed us to separate the recording into two intervals and compare the MU velocity twitch profiles of a unit active throughout the whole recording with and without the second MU active. Fig. 8 shows that the STA peak-to-peak amplitude decreased when a nearby unit was active for an MU active throughout the whole recording time (see Supplementary materials for further examples). Therefore, the activity of an MU is affected by its neighbors. Interestingly, the STA of a motor unit never increased in amplitude when a second unit became active (it always decreased), ruling out the possibility that the effect was due to synchronized MU activity, which would have conversely increased the STA amplitude [26]. This supports the results from experiment 1 – once the nearby unit is active, the already active unit will contribute with a different velocity twitch that, on average, is of lower amplitude.

**Fig. 8.**
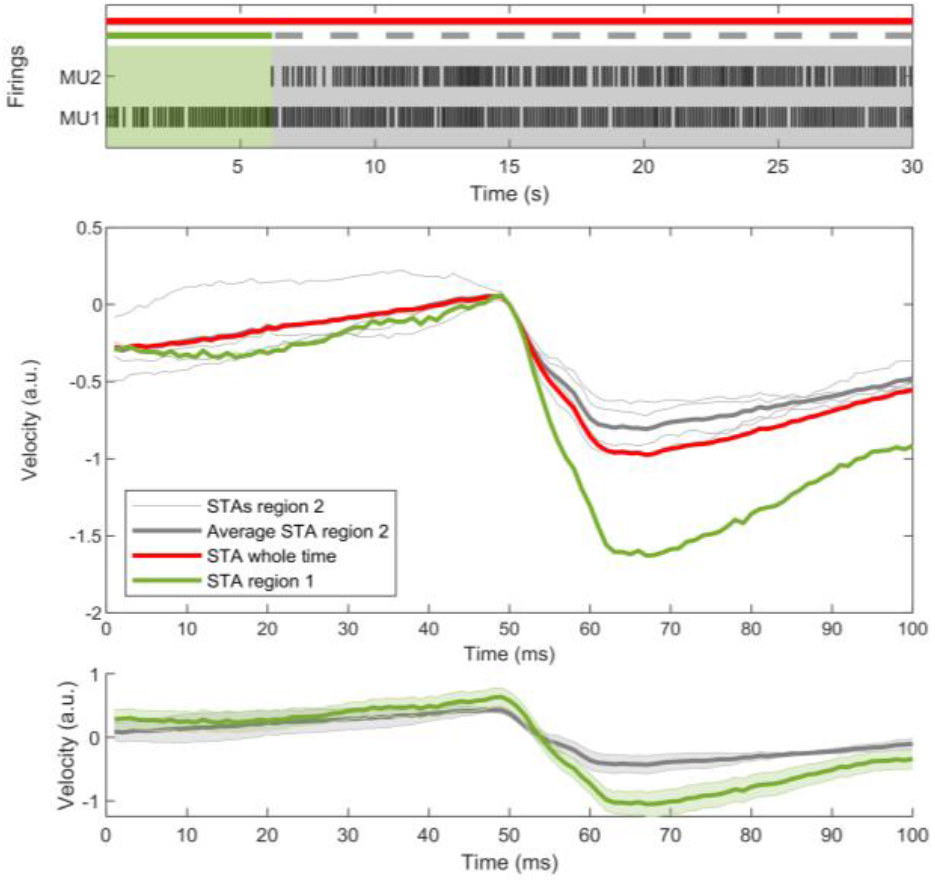
Example spike-triggered averages (STA) for a motor unit (MU1) before and after the activity of a nearby motor unit (MU2) has started. Red curve is the STA for MU1 over the whole time (n = 245). Green curve is STA of MU1 using all spikes in the green region of the upper plot (n = 46). Thin grey lines are STAs with randomly distributed spikes of MU1 from the grey region of the upper plot, ensuring equal numbers of triggers as used for green curves (n = 46). The thick grey curve is the average of the thin grey curves (n = 4). In the middle plot all curves are set to 0 at 50 ms (time of firing) for ease of comparison. The peak-to-peak amplitude of MU1 STA decreases when MU2 is active – the activity of MU2 suppresses the activity of MU1. The lower plot shows the green and grey curves with their standard errors.

For the example of Fig. 8 (and those in the Supplementary materials), there is a mean increase in discharge rate between green and grey segments, which could increase fusion and cause a decrease in velocity fluctuations. However, this change is of approximately 0.5 pulses per second and could only, therefore, explain a very small change in magnitude (maximum decrease of approximately 4% [27], [28]). In contrast, we saw an average decrease of 45% in peak-to-peak amplitude. Hence, a change in fusion cannot explain the decrease in amplitude.

Fig. 9 shows an example of the analysis (ii) described in section I.B.4 (see Supplementary materials for a further example). The middle plots show that the STA using the ‘unaligned’ firings only differs from that using the ‘aligned’ firings for one unit (MU1 in A), suggesting a non-symmetrical relation between units. From the bottom plot, we observe that the relation between velocity twitch shape and time to the closest firing of the other unit is complicated – this is likely to depend on geometry: overlapping MUs with aligned velocity twitches may reinforce each other, whereas neighboring units with overlapping velocity twitches may oppose each other and cancel out. Furthermore, because of the long duration of the velocity twitches, the ‘unaligned’ group also includes the activity of the two units partly overlapped in time.

**Fig. 9.**
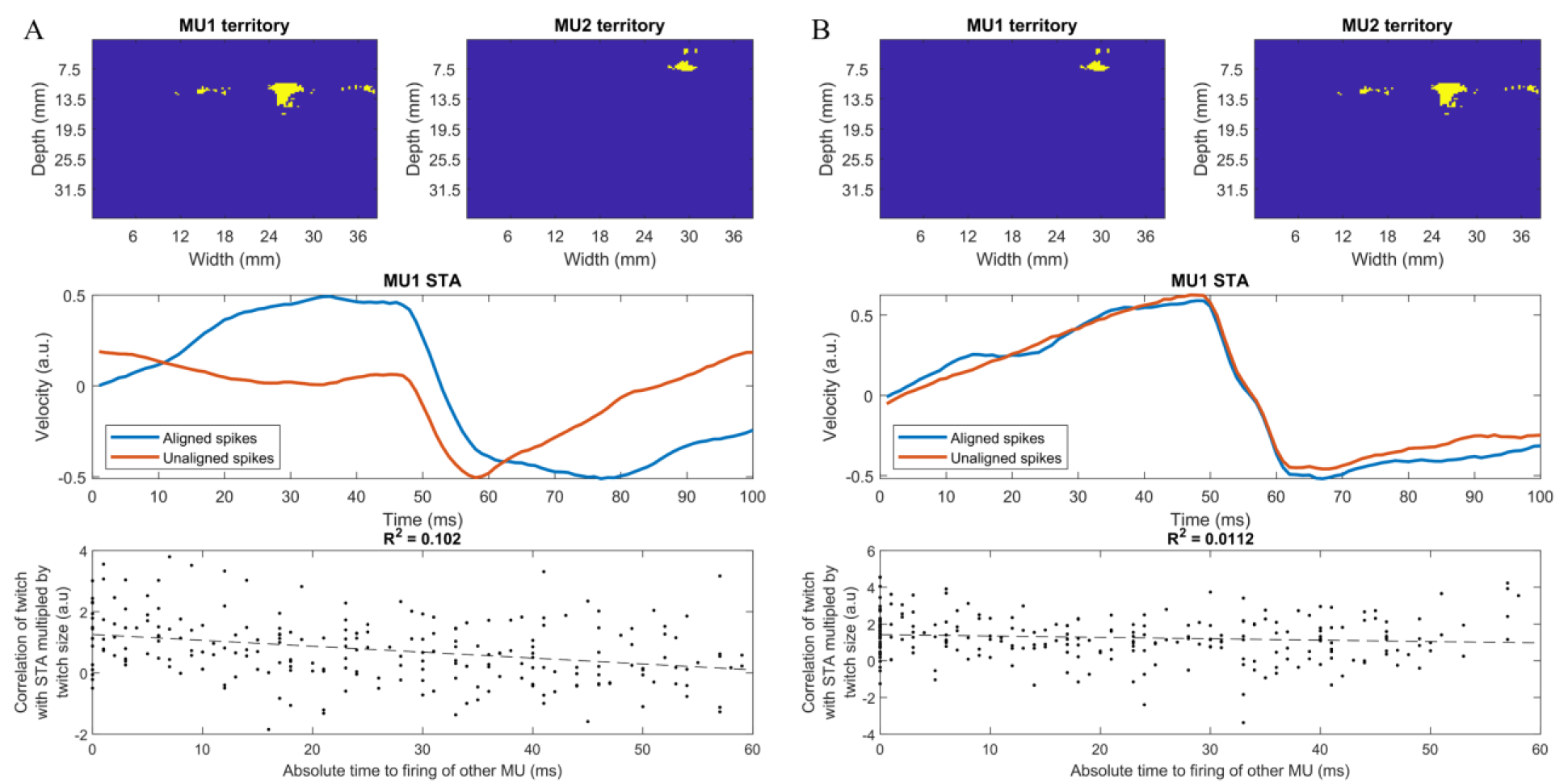
A: Top panel: regions of MU1 and MU2. Middle panel: The blue curve is an STA of MU1 with firings from MU1, which are within a set window around firing times of MU2 (window adjusted in each example to ensure an equal number of firings used for each STA). The red curve is an STA of MU1 with firings from MU1 which are not within the set window around the firing times of MU2. The difference in these curves shows that the proximity of a firing of the nearby unit impacts the twitch of a given unit. Bottom panel: for each firing time of MU1, the correlation of the twitch with the overall STA multiplied by the size of the twitch is plotted against the absolute time to the closest firing of MU2. A very weak linear relationship is shown (R2 = 0.102). B: The motor units used in A are swapped. In the middle panels, no difference is seen between the blue and red curves, suggesting that, in these cases, the firing of the nearby unit does not affect the unit’s twitch profile. The bottom panels show an approximately 10 times weaker linear relationship (R2 = 0.0112). Although a MU is affected by the firing of another nearby unit, this relationship is not reciprocal, and thus the other unit may not be impacted.

## IV. DISCUSSION

We used concurrent HDsEMG and US, and iEMG and US recordings to analyze the linearity of velocity twitch summation with respect to local muscle velocity in voluntary contractions. Using state-of-the-art HDsEMG MU decomposition, we compared STA twitches for one MU across increasing contraction forces, thus with increasing co-active units. We hypothesized that if the system is linear, the effects of the activation of neighboring units (producing twitches at random times relative to the studied unit) will not change the average velocity profile obtained using the STA process, resulting in unchanging average twitch profiles at increasing force levels. Conversely, we found that the peak-to-peak amplitude decreased. Thus, we conclude that there are mild non-linearities present in the system. We believe the non-linearity results in a non-stationary velocity twitch that changes depending on the statistics and times of discharge of other nearby MUs, because of the mechanical coupling of the MUs.

The straightforward way to test for linearity would have been to activate two MUs separately and concurrently and compare their activity in the two cases. However, this approach has practical constraints in voluntary contractions. First, MUs are recruited sequentially, therefore, individually activating the higher-threshold MU is often impossible. Furthermore, stable individual MU control over long time intervals may be challenging for naїve subjects [2]. Instead, we concurrently recorded HDsEMG and US at 4 stable contraction forces. Since the number of active MUs increases with increasing force, the different forces correspond to different numbers of active MUs [29]. In these conditions, if the velocity fields of MUs are independent and linearly superimpose, increasing the number of active MUs will not impact the velocity twitches. Using the STA method, we therefore compared the velocity twitches by tracking MUs across forces to determine the system linearity.

Contrary to global muscle force which is the combination of all MU outputs into a one-dimensional temporal signal (summing all activity over space), the US velocity signals have both spatial and temporal dimensions, increasing the sparseness of the system. Hence, we can observe velocity twitches that are isolated from the twitches of other units. We assumed that the isolated twitches were those of more consistent shape over time, and we identified them as the ‘correlated’ twitches. The correlated velocity twitches did not change in amplitude when the number of active MUs increased, indicating that the changes in the grand average twitch are due to temporally local non-linearities that occur when two or more MUs discharge close in time and space. This also excludes that the observed non-linearities are due to a change in the whole muscle properties, at least in the range of investigated forces.

In all data shown here, the MU velocity twitch was negative (here defined as away from the US probe). However, in many cases, we observed examples of regions that also moved toward the probe. These may be in conjunction with negative motion (MU twisting/complicated motion), or the only motion associated with that unit. Detection of motion away from the probe is easier as any motion towards the skin may be reduced due to compression from the probe or may result in small skin motions and hence of the probe, thus minimizing relative motion. As such, the data in this work are limited to motions away from the probe. However, the same conclusions can be drawn when considering the positive motions (results not shown).

Understanding the workings of intra-muscular motion has broader implications than US interpretation and processing, such as in musculoskeletal modelling. Generally, a muscle is modeled as a single actuator [30], geometrically described as a line segment between its insertion points on the skeletal bodies [31], assuming all fibers have the same length and neuromuscular properties using multiscale simplifications [32]. Some muscle modeling developments have aimed for a more physiological underpinning by describing the dynamics of the individual MUs of the muscle, for example by using controlled pools of in-parallel mathematical models of individual MUs with artificial [29], [33]–[35] and experimentally generated [36] MU firing times. Other studies have developed volumetric representations of muscles described as collection of spatially-arranged fibers [37], [38]. However, both approaches assume no inter-unit connectivity between the modeled fiber actuators. In such cases, the relative motions between fibers are assumed to be independent and their generated forces are modelled to add linearly, which contradicts physiological findings [32], [39]. Experimental insights into the true impacts of the inter-relation between MUs could improve these models, and US detection methods could enable a precise mapping of MUs within the muscle volume.

The linearity of twitch summation for multiple active MUs has previously been considered from two main perspectives: the skin displacement via mechanomyography (MMG) and the force output at the tendon. Using the former method, Orizio et al. [40] found that at very low stimulation frequencies MMG twitch summation can be considered linear, however within the physiological range of frequencies, MMG twitch summation is non-linear, resulting in a different MMG signal to the algebraic sum of the individual signals. However, MMG is measured at the skin surface. Thus, it does not provide a local measure of muscle displacement.

Studies utilizing force measurements have had a large variety of results, mainly showing non-linear force twitch summation. In the cat medial gastrocnemius and cat soleus [41], [42] the simultaneous activation of multiple MUs resulted in a larger force output than the algebraic sum of the individual MUs. In contrast, the cat soleus [42] and tibialis anterior [43] muscles showed lower force outputs when simultaneously active than their algebraic sum. The rat soleus [44] showed almost linear summation, whereas the rat gastrocnemius [45] and the cat peroneus longus [46] showed an MU-dependent mix of super-additive and sub-additive results. Results are highly variable for different muscles and species, likely resulting from the large differences in fiber orientation, pennation, length, connectivity, and force transmission to the tendon. The non-linearity observed in force studies is attributed to muscle fiber connectivity and pennation angle. The present study shows that local muscle movements do not sum linearly, which is consistent with the interpretation of non-linearity as attributed to intra-muscular mechanical interaction.

Using iEMG, we could study MUs with very close territories because of the small detection volume of the iEMG electrodes. Here, we could directly study cases where MU activity was initiated mid-way during the contraction. In all cases, the velocity twitch of a given MU was suppressed when the activity of a nearby unit began. This supports the conclusions of experiment 1 – the nearby units’ activity impacts the velocity twitch profile of a given unit and causes the velocity twitch profile to decrease in amplitude. However, a new finding from experiment 2 (Fig. 9) was that MUs do not influence each other in symmetrical ways. When looking at pairs of MUs, one unit appears influenced by the firing of the other more than vice-versa.

Fig. 9 shows the non-symmetrical effect of MUs on their velocity twitches. It is reasonable to assume that a larger unit would have a greater effect on the velocity field of the muscle with respect to a smaller unit, as it involves the movement of more fibers. According to the size principle [47], [48] the later recruited units will be larger, so we would expect that all the ‘MU2’s in Fig. 8 are larger than the ‘MU1’s, albeit the difference would be small as the recruitment thresholds are very close together. In each case here, we see that the firing of MU2 affects MU1, as we expect from the recruitment order reasoning. However, we have not investigated the effect of MU1 on MU2. Further protocol development to allow for investigation into recruitment order and MU velocity twitch profiles should be done using iEMG and US on a larger MU population.

## V. CONCLUSION

We used ultrafast US in conjunction with both HDsEMG and iEMG to study the linearity of the summation of velocity twitches of active MUs. We found that at increasing force levels, where increasing numbers of MUs are active, the STA twitch amplitude monotonically decreased. Hence the system is non-linear. We conclude that linear methods for decomposition are unlikely to successfully decompose full velocity twitch trains from US image series. However, when considering only a subset of discharge times, we were able to extract a velocity profile not influenced by force. This result indicates that the non-linearity of the system is not due to the properties of the whole muscle but rather to local mechanical coupling between muscle units. Finally, we observed that the influence of one MU on another MU velocity profile is not symmetrical. These results provide information to design more advanced US decomposition algorithms than currently available and to interpret the contribution of individual MUs to muscle movement.

## ACKNOWLEDGMENT

JI was supported by a Ramón y Cajal grant (RYC2021-031905-I) funded by MCIN/AEI/10.13039/501100011033 and UE’s NextGenerationEU/PRTR funds. RR was supported by Hans Werthén Foundation and the Swedish Research Council for Sport Science (D2023-0003).

