## Supplementary Material for "Non-linearity in motor unit velocity twitch dynamics: Implications for ultrafast ultrasound source separation"

### Supplementary materials

Fig 1 and Fig. 2 show further examples from experiment 2, as discussed in section III.B.

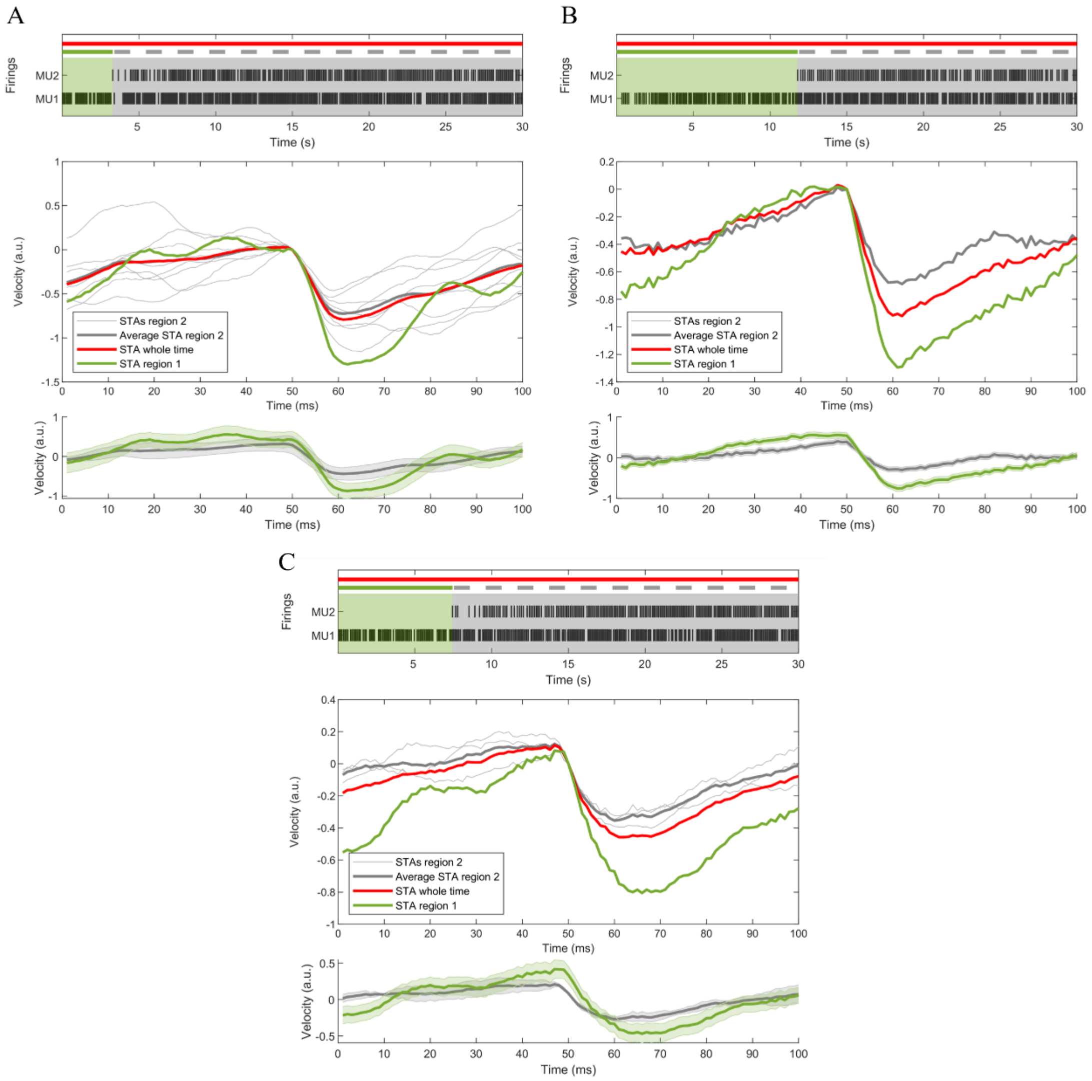


Fig. 1. Examples of spike-triggered averages (STA) for a motor unit (MU1) before and after the activity of a nearby motor unit (MU2) has started. Red curves are STAs for MU1 over the whole time (n = 311, 285, 288 respectively for A, B, C). Green curves are STAs of MU1 using all spikes in the green region of the upper plot (n = 34, 109, 69). Thin grey lines are STAs with randomly distributed spikes of MU1 from the grey region of the upper plot, ensuring equal numbers of triggers as used for green curves (n = 34, 109, 69). Thick grey curves are averages of thin grey curves (n = 8, 1, 3). All curves are set to 0 at 50 ms (time of firing) for ease of comparison. In every example, the STA twitch velocity profile is greater in the green region than in the grey region. The activity of MU2 suppresses the activity of MU1. The bottom plot in each case shows the curves with standard error.

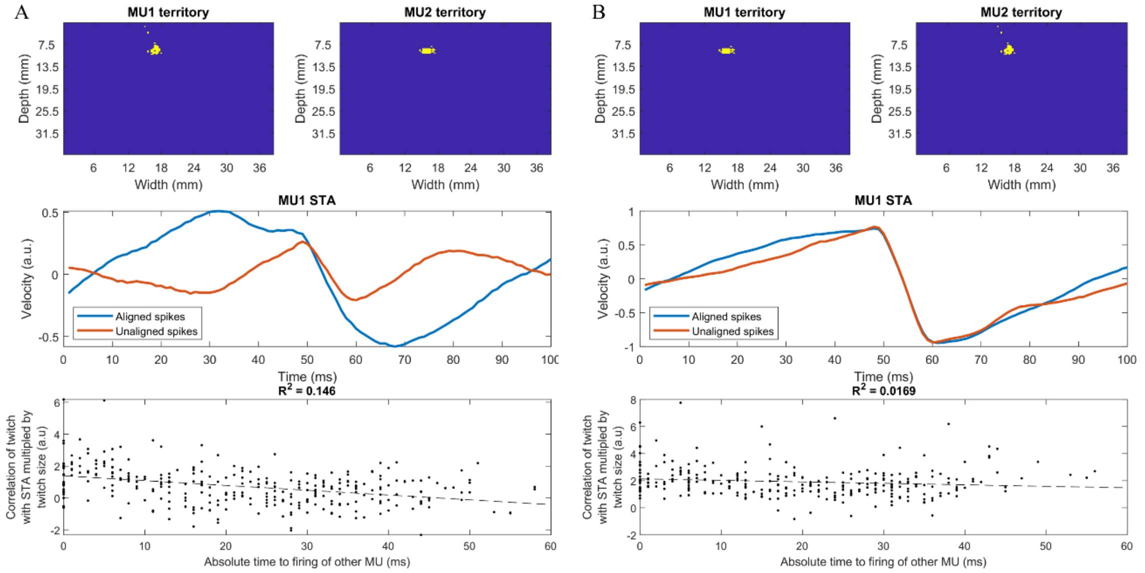


Fig. 2. A: Top panel: Regions of MU1 and MU2. Middle panel: The blue curve is an STA of MU1 with firings from MU1, which are within a set window around firing times of MU2 (window adjusted in each example to ensure an equal number of firings used for each STA). The red curve is an STA of MU1 with firings from MU1 which are not within the set window around the firing times of MU2. The difference in these curves show that the proximity of a firing of the nearby unit impacts the twitch of a given unit. Bottom panel: For each firing time of MU1 the correlation of the twitch with the overall STA multiplied by the size of the twitch is plotted against the absolute time to the closest firing of MU2. A very weak linear relationship is shown ($R^2$ = 0.146). B: The motor units used in A are swapped. In the middle panels, no difference is seen between the blue and red curves, suggesting that in these cases the firing of the nearby unit does not affect the unit's twitch profile. The bottom panels show an approximately 10 times weaker linear relationship ($R^2$ = 0.0159). Although a MU is affected by the firing of another nearby unit, this relationship is not reciprocal, and thus the other unit may not be impacted.
